# Transcranial Direct Current Stimulation Alters the Waveform Shape of Cortical Gamma Oscillations

**DOI:** 10.1101/2022.04.25.489371

**Authors:** Tom R. Marshall, Andrew J. Quinn, Ole Jensen, Til Ole Bergmann

**Affiliations:** Wellcome Centre for Integrative Neuroimaging, Department of Experimental Psychology, University of Oxford, United Kingdom; Oxford Centre for Human Brain Activity, Wellcome Centre for Integrative Neuroimaging, Department of Psychiatry, University of Oxford, United Kingdom; Centre for Human Brain Health, School of Psychology, University of Birmingham, United Kingdom; Neuroimaging Center (NIC), Johannes-Gutenberg University Medical Center Mainz, Germany; Leibniz Institute for Resilience Research (LIR), Mainz, Germany

## Abstract

Neuronal oscillations in different frequency bands have been linked to a wide variety of cognitive functions, and may even be a fundamental mechanism of inter-regional communication. For this reason, *manipulation* of oscillatory activity via brain stimulation is a central goal in neuroscience research. However, the vast majority of studies characterise oscillatory activity solely in terms of *amplitude* and *frequency*. Oscillations can also be characterised by their *waveform shape*; the degree to which they resemble or deviate from sinusoids. Here we exploit Empirical Mode Decomposition (EMD), a novel method that allows quantification of oscillatory waveform shape. We show for the first time that transcranial direct current stimulation (tDCS) alters the waveform shape of gamma oscillatory activity in the visual cortex. Notably, changes in waveform shape were limited to one half of the phase cycle; anodal stimulation led to a relatively slower, and cathodal to a relatively faster, descending half-wave. tDCS is generally believed to affect cortical excitability via alteration of resting membrane potential. Interestingly, simulations of altered cortical excitability in a gamma-generating neuronal population indicated the waveform shape changes observed experimentally likely stem from stimulation of pyramidal neurons. These findings have implications for understanding the neural consequences of tDCS at the level of neuronal population phenomena such as cortical oscillations and underscore the importance of waveform shape as an important feature of neuronal oscillations.

## Introduction

A central goal in cognitive neuroscience is to use transcranial brain stimulation techniques to alter brain activity, and thereby cognitive function^1^. One promising candidate is transcranial direct current stimulation (tDCS)^2^: the application of weak electric currents to the scalp to produce localised changes in functional properties of neurons. tDCS is safe, affordable, non-invasive, and has been shown to be effective at altering cortical excitability, with effects on cognition and behaviour^3–5^.

However, behavioural consequences of tDCS have been characterised as weak or inconsistent^6,7^. This is unsurprising considering its putative mechanism of action. tDCS is generally accepted to alter neuronal excitability^8^, however, both the relationships between neuronal excitability and neuronal population activity^9^, and between population activity and cognitive processing^10^, are extremely complex. To build better bridges to behaviour, it is therefore important to understand the effects of tDCS at the level of neuronal population signals, such as neuronal oscillations.

Rhythmic neural synchronization within and between brain regions is known to subserve a wide variety of cognitive functions, and synchronisation in the high-frequency gamma range may be a general mechanism for inter-neuronal communication^11^. Changes in cortical gamma activity have been linked to attention^12,13^, memory^14^, and learning^15^. Therefore, understanding the possible effects of tDCS on gamma-band activity could be a fruitful avenue to optimise its effects on cognition and behaviour.

Theoretical models of gamma oscillations tend to posit alternating periods of excitation and inhibition^16^, with key features of the activity determined by biophysical properties. For example, the duration of inhibitory synaptic currents is a strong determinant of frequency; the faster excitatory neurons can recover from inhibition and fire again, the shorter the overall period of the oscillation^17,18^. Speculatively, one might therefore expect a cortical excitability change to have different effects on different phases of the gamma oscillation; e.g., shortening the inhibitory phase without altering the excitatory phase, or *vice versa*. In other words, changing cortical excitability might alter the *waveform shape* of gamma oscillations.

Recently attempts have been made to manipulate gamma-band activity by application of tDCS to the scalp. The majority of studies have examined so-called *offline effects*; comparing gamma activity before and after tDCS application^19,20^. Methodological advances^21^ have enabled the measurement of *online* tDCS effects on gamma oscillations^22,23^; simultaneously stimulating the brain and recording whole-brain activity with magnetoencephalography (MEG).

However, the majority of studies of visual gamma in humans have tended to quantify the *amplitude* and *frequency* of the gamma oscillations. While both gamma amplitude^24^ and frequency^25^ have been shown to be important for stimulus processing, it is possible that tDCS – particularly *online* tDCS – affects gamma-generating neuronal populations in ways that are not easily detected in these features of the signal.

Here, we reanalysed a concurrent tDCS/MEG dataset where participants viewed stimuli known to generate strong visual-cortical gamma oscillations during whole-brain MEG recording, while visual cortex was concurrently stimulated with tDCS^23^. We extracted gamma oscillations on a single-cycle basis, and exploited recent methodological advances^26,27^ to quantify their amplitude, frequency, and waveform shape under different tDCS conditions (anodal, cathodal, sham).

Importantly, gamma oscillations are a population-level phenomenon that depend on complex interactions between diverse cell types^17^. Furthermore, tDCS is believed to differentially affect different cell populations due to their distinct morphologies^28^. Since MEG cannot directly provide insights into changes at the neuronal level, we validated our experimental results using a biophysical model of gamma oscillations comprising simulated pyramidal and interneurons, where we simulated tDCS by changing the excitability of the neuronal populations.

## Methods

These data were originally reported in^23^. 20 participants (10 female, mean age 24.5 years) took part in the study, with one participant excluded due to falling asleep during data acquisition. All participants gave informed consent and the study was approved by the CMO region Arnhem-Nijmegen (2014/138).

### Experimental task

The experimental paradigm (Fig. 1A) was previously described in^23^. Briefly, after fixating a foveal dot for a 1400ms baseline period participants observed an inwardly moving annular grating (spatial frequency 2.5 cycles/dva, size 8 dva, speed 0.8 dva/second, contrast 40%) for 1400ms, and were instructed to press a button with their right index finger when the stimulus increased in speed (to speed 1.12 dva/second, 1/6 trials). Speed-change detection was near ceiling for all participants and speed-change trials were not analysed; their purpose was to ensure stable attention to the stimulus.

**Figure 1:**
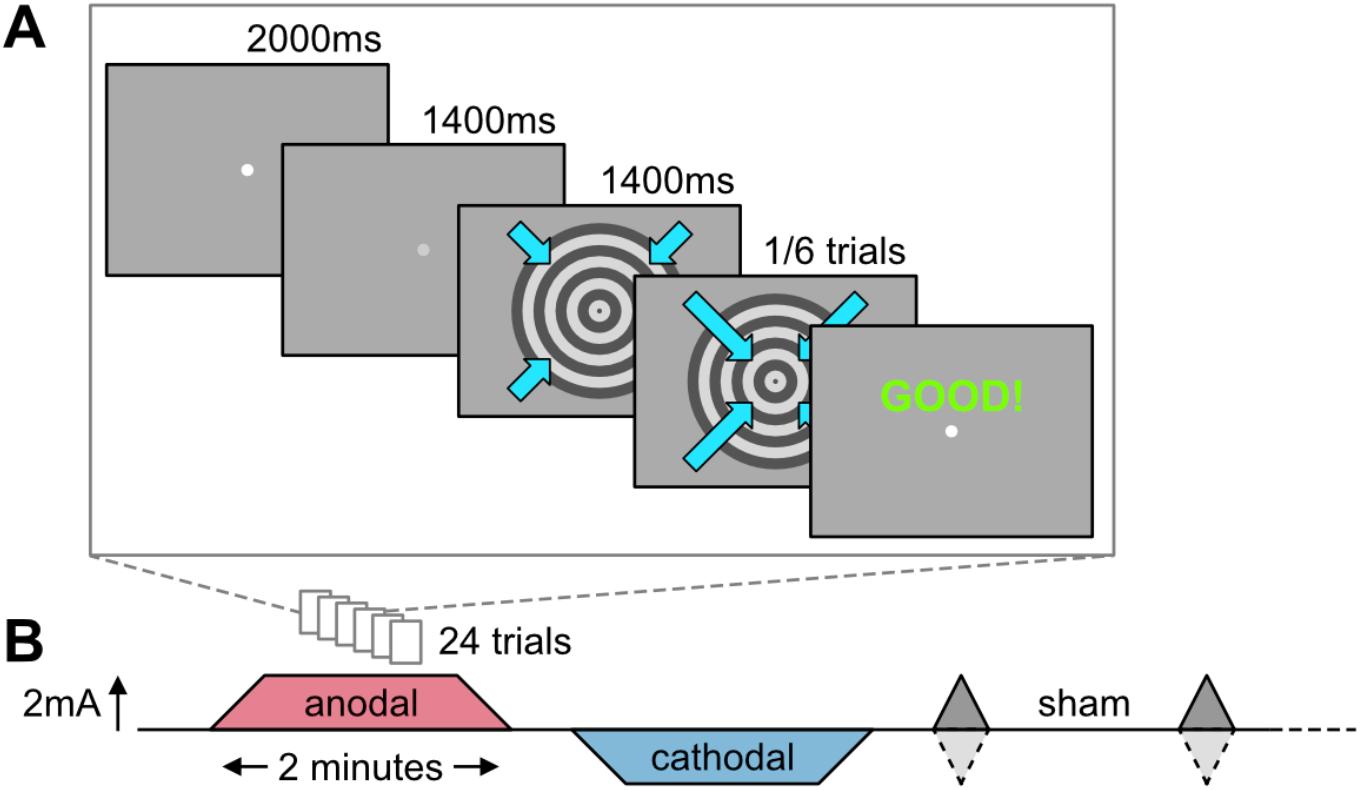
Task and tDCS parameters. A) Structure of a single trial. B) Description of blocked experimental design (interleaved blocks of anodal, cathodal and sham stimulation) and tDCS parameters across the experiment.

### tDCS

While participants performed the task tDCS was applied to the scalp in a Cz/Oz montage (Cz electrode size 5 × 5 cm = 25cm^2^, Oz electrode size 10 × 5 cm = 50cm^2^). tDCS was applied in 15 2-minute blocks; 5 blocks of 2mA anodal stimulation and 5 blocks of 2mA cathodal stimulation, ramping up and down over 5 seconds, and 5 blocks of sham stimulation, ramping up to 2mA and immediately down again over 5 seconds at the start and end of each block (Fig. 1B). Blocks were interleaved so no two active blocks of the same polarity occurred in succession to prevent build-up of tDCS aftereffects. During each block participants completed 24 trials. Trials occurring during tDCS ramping were not analysed due to the presence of strong ramping artifacts in the MEG signal. Trials during active tDCS contained low-frequency artifacts time-locked to the heartbeat and possibly arising from the pulsation of cranial blood vessels causing small movements of the stimulation electrodes; these artifacts were attenuated but not removed by beamforming^23^.

### MEG preprocessing

MEG preprocessing and analysis was performed in a combination of MATLAB (version 2021a, http://www.mathworks.com) and Python (version 3.5.4, http://www.python.org). Preprocessing has already been described in^23^. Briefly, a two-step beamforming approach was applied: Firstly, we used frequency-domain DICS beamforming^29^ to compare activity in the pre-stimulus and post-stimulus periods in the broad gamma band (45-75 Hz) at several thousand cortical locations, and selected the voxel for each participant showing the largest gamma-band power increase. In all 19 participants this voxel was in occipital cortex near the midline. Power increases ranged from 48% to 336% (mean 144% standard deviation 76%). Secondly, we used time-domain LCMV beamforming^30^ to reconstruct the time course of activity at the selected peak-gamma voxel. Sensor covariances were computed for each 4-second epoch for all trials, pooling across tDCS condition, from which a common spatial filter was derived for the selected voxel. Raw MEG data were then multiplied with this common filter to produce a single ‘virtual channel’ time series. Note that the resultant time series is inherently sign-ambiguous; a positive deflection is equally likely to correspond to the ‘peak’ or the ‘trough’ of an oscillation. We used a group PCA approach to resolve this sign ambiguity across participants; see section ‘Resolving dipole sign ambiguity’.

### Empirical mode decomposition

Conventional analysis of oscillatory activity relies on the Fourier transform, which rests on the proof that arbitrary time series can be decomposed into a set of sinusoidal basis functions. While hugely powerful, this approach creates challenges for understanding non-sinusoidal features of brain signals^31^. Therefore we instead used Empirical Mode Decomposition, an alternative approach that separates data into different components (‘intrinsic mode functions’ or IMFs) containing progressively slower neural dynamics.

Empirical mode decomposition (EMD) was performed using the EMD Python toolbox^27^ (version 0.3, https://pypi.org/project/emd/). Epoched virtual channels were concatenated, then decomposed into a set of five intrinsic mode functions (IMFs). Sifting was performed using a set of dyadic masks (‘mask sift’); sine-waves decreasing by a factor of two. This has been shown to reduce mixing of different frequencies into a single IMF. The frequency of the first mask was derived from the instantaneous frequency of the first IMF after a standard, non-masked sift of the data. Mask amplitude was computed as a ratio of the standard deviation of the input signal. Signal envelopes were interpolated using Piecewise Cubic Hermite Interpolating Polynomial (pchip) interpolation. The decision to stop at five IMFs was arbitrary, however time-frequency analysis of the IMFs showed that relevant and expected oscillatory responses (visual stimulus-induced gamma-band increase, alpha and beta-band decrease) were captured in the first five IMFs (see Fig. 3). Note that, as EMD is sequential, estimating further IMFs would have no effect on the IMFs already estimated.

To ensure well-established task components (stimulus-induced gamma band increase, alpha/beta decrease) were captured by the IMFs we performed time-frequency analysis on each IMF using the Fieldtrip MATLAB toolbox^32^ (https://www.fieldtriptoolbox.org/). For low frequencies, we computed power estimates at integer frequencies from 1-30Hz using a 300ms sliding time window moving over the data in 50ms steps and multiplied with a Hanning taper. For high frequencies, power at frequencies between 30 and 150Hz (4Hz spacing) was estimated using a set of 3 orthogonal Slepian tapers applied to a 500ms sliding time window in 50ms steps.

### Resolving dipole sign ambiguity

Two hypothetical current dipoles with opposite polarities and opposite orientations will generate identical fields at the MEG sensors. For this reason, source-reconstruction methods such as beamforming that perform the *inverse* operation – infer the time course of a dipole from the fields measured at the sensors – cannot distinguish the polarity of a source; the recovered source-space time courses are inherently sign-ambiguous. This is problematic for group analysis of waveform shape; sign ambiguity means that ‘peaks’ found in the time courses of one participant may correspond to ‘troughs’ in another, and vice versa. It is therefore necessary, prior to waveform shape analysis, to resolve the sign ambiguity by ‘aligning’ participants’ data based on some data feature. Based on the assumption that the gamma intrinsic mode function captured the activity of the underlying gamma source generator, we aligned data across participants using the timelocked stimulus-evoked response of this IMF.

For each participant we averaged over all trials and cut out a 550ms window around visual stimulus onset (100ms pre to 450ms post). A clear event-related response was visible from 100-200ms after stimulus onset (Fig. 2A, upper), which appeared inconsistent across participants, and therefore highly attenuated in the grand average (Fig. 2A, lower, blue). To resolve the sign ambiguity we created a matrix with each row representing one participant and each column one time point, performed a PCA on this matrix, and took the sign of the first principal component for each participant. In participants where the sign was negative, we sign-flipped their data. This produced event-related responses that were clearly strongly aligned and highly similar across participants (Fig. 2B). We then sign-flipped the raw data in the same participants and carried out all subsequent analysis on the aligned data. Note that this is not considered double-dipping for two reasons: firstly, the flip decision was based on the phase-locked ‘evoked’ responses shortly after stimulus onset, while our main analyses focused on the waveform shape of non-phase-locked ‘induced’ gamma oscillations in the later ‘sustained gamma’ period. Secondly, our critical comparisons concerned differences between tDCS conditions (anodal, cathodal, sham), whereas this processing step was based on the event-related average collapsing across all three conditions.

**Figure 2:**
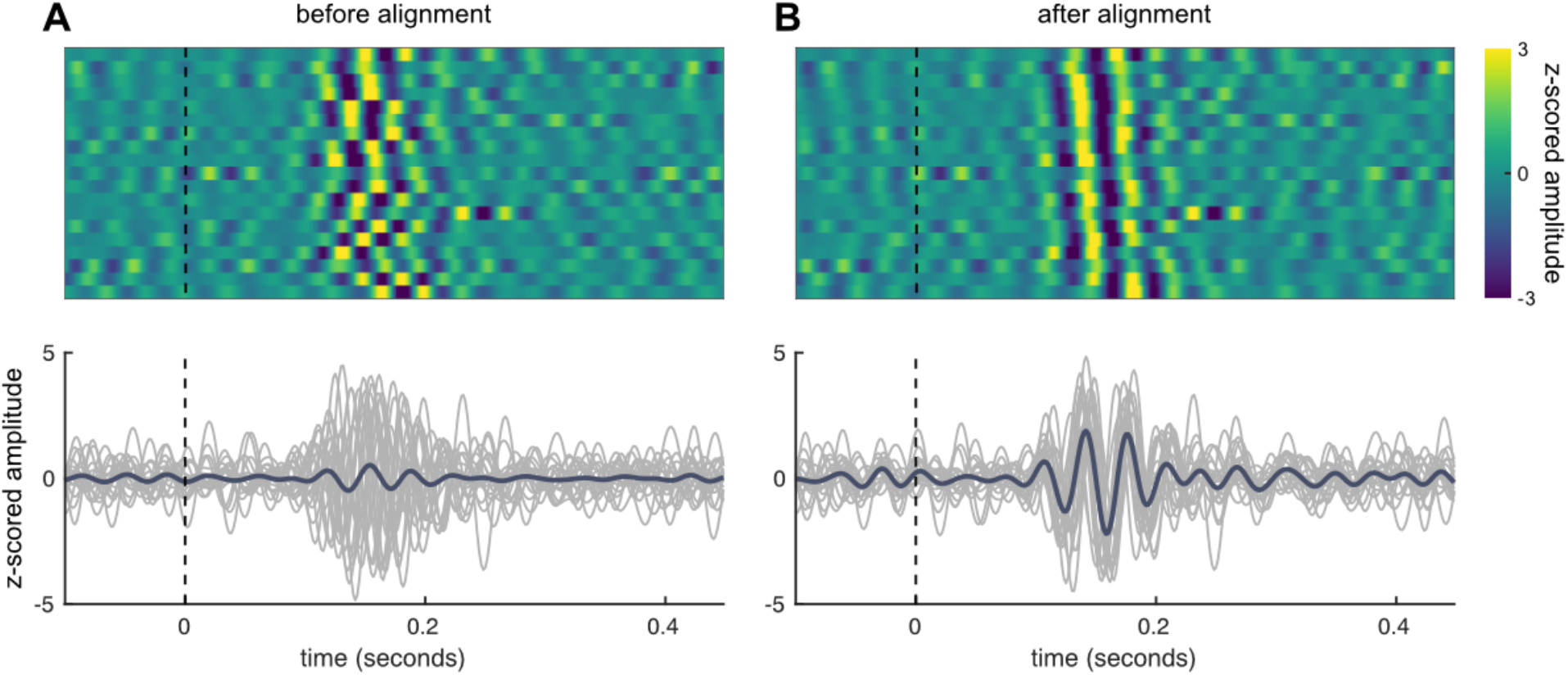
PCA-based alignment to resolve dipole sign ambiguity. A) Upper, z-scored averaged event-related fields from gamma IMF. Lower, event-related averages for individual participants (grey) and group average (blue). B) As A, but after PCA-based realignment.

**Figure 3:**
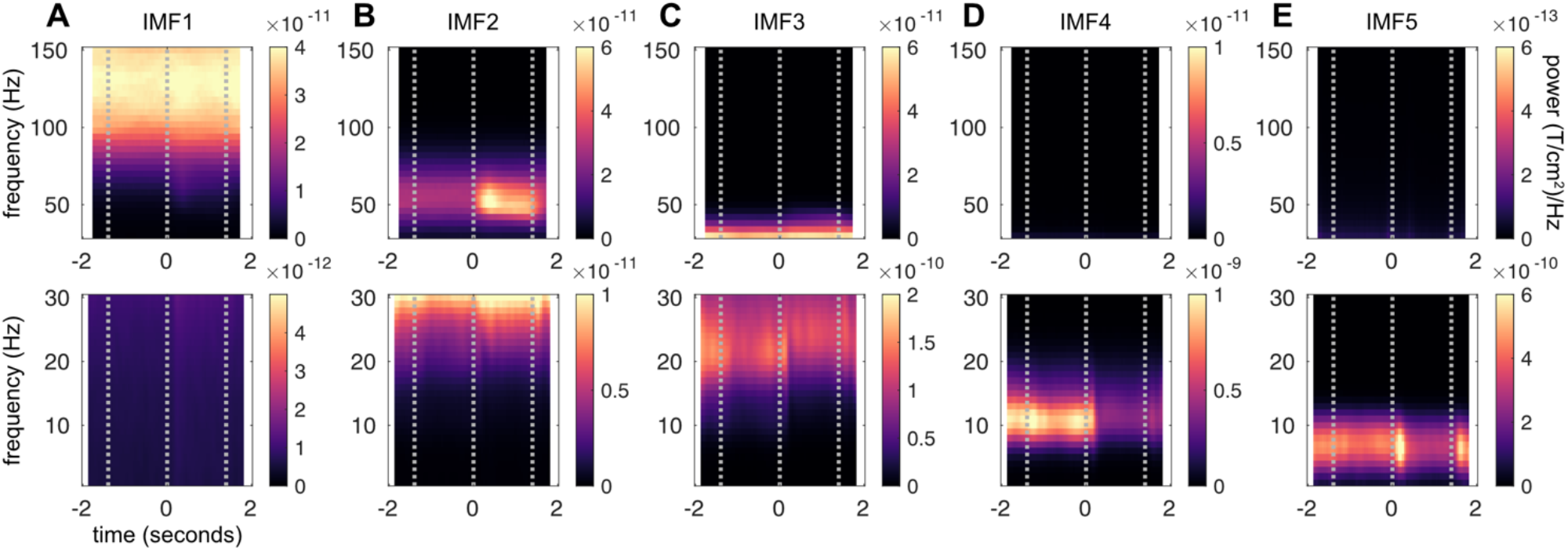
Time-frequency analysis of intrinsic mode functions (IMFs). A-E) Fourier analysis of intrinsic mode functions resulting from EMD on beamformed MEG data. IMFs capture oscillatory activity in progressively lower frequency bands, as well as well-understood task features (B, gamma-band power increase, C-E, alpha/beta-band power decrease). Note: Raw power is shown, rather than baseline-corrected power changes, to illustrate the absence of oscillatory power at certain frequencies throughout the epoch. Grey lines indicate, from left to right: Fixation dot on (beginning of baseline period); visual stimulus on; visual stimulus off.

### Cycle-by-cycle analysis

Individual cycles were detected in the gamma-band IMF (*emd*.*cycles*.*get_cycle_inds*). Instantaneous phase, frequency and amplitude of each mode were extracted via a normalised Hilbert transform. The instantaneous phase values were then used to define individual cycles according to the criteria: strictly positively increasing phase, beginning at zero and ending at 2*pi and containing 4 unique ‘control points’; ascending zero crossing, peak, descending zero crossing, trough^26^. Having isolated individual ‘good’ cycles, summary stats – time of occurrence, duration, mean amplitude, and mean frequency – were calculated for each cycle. We then further classified cycles according to their latency within a trial, and the tDCS condition (anodal, cathodal, sham). For each participant we then averaged across all trials in each tDCS condition to calculate mean cycle amplitudes and cycle frequencies at every time point. We detected and excluded spurious cycles at the trial boundaries arising from the concatenation of epoched data. This accounted for 0.5% of all cycles.

Since we knew the moving visual stimulus produced strong changes in gamma amplitude, we tested whether this effect was present when analysing the gamma IMF at the single-cycle level. Accordingly, we extracted cycle info from a pre-stimulus baseline window of -1.1 seconds to -0.3 seconds, and a ‘visual stimulus on’ window from 0.3 to 1.1 seconds i.e., the entire baseline and stimulus-on windows minus the very beginning and very end, to avoid contamination by stimulus on- and offset. We then averaged over epochs (baseline, stimulus on) and tDCS conditions (anodal, cathodal, sham), and compared gamma cycle amplitude between baseline and trial using paired t-tests, and between tDCS conditions using within-participant ANOVA and post-hoc paired t-tests.

### Waveform shape analysis

A full description and rationale for describing waveform shape in terms of phase-aligned instantaneous frequency can be found in^26^, and a brief sketch can be found in supplementary materials (‘the logic of phase alignment’). Waveform shapes for each detected cycle were calculated by estimating the instantaneous phase (IP) from the Hilbert transform, taking the temporal derivative of IP to obtain instantaneous frequency, then ‘phase aligning’ all cycles by interpolating the instantaneous frequency values to a common ‘phase space’. We took phase-aligned instantaneous frequency for each cycle within the ‘visual stimulus on’ window (+0.3 to +1.1 seconds) and computed average waveforms for each participant and tDCS condition (anodal, cathodal, sham). To statistically compare tDCS conditions we used cluster-based permutation tests^33^. Briefly; t-values comparing anodal and cathodal tDCS were calculated, from which the ‘max-summed cluster’ (greatest sum of contiguous significant t-values) was computed. The labels (anodal, cathodal) were then permuted 5000 times to create a reference distribution of 5000 cluster-t-values under the null hypothesis, with the true cluster statistic evaluated against this reference distribution. Since clustering is typically performed over non-wrapping axes (time, frequency, space), whereas phase is a wrapping axis, we adapted the clustering algorithm slightly to allow clusters to be formed across the cycle boundary (e.g., comprising the last, first and second t-values). For illustrative purposes comparisons against sham were performed in the window of significant anodal/cathodal difference.

### Biophysical modelling

To model gamma oscillations in visual cortex we used Human Neocortical Neurosolver^34^ (HNN); a configurable neocortical circuit model that simulates currents measured by MEG and EEG. We adapted the standard circuit model of gamma oscillations (https://jonescompneurolab.github.io/hnn-tutorials/gamma/gamma) whereby a Poisson input to the simulated region produces a narrow-band rhythmic activity in the gamma range. The standard ‘weak ping’ model implemented in HNN has some features that fall short of biophysical plausibility. For example, the neuronal populations in layer 2/3 and layer 5 are not connected to each other, essentially producing two independent gamma sources. While laminar differences and intralaminar interactions in gamma activity are of interest (for example^35–37^) we considered this beyond the scope of the present study. Therefore, we reduced the tonic drive to L2/3 to zero and focused only on the L5 signal, which – due to longer apical dendrites – is the primary contributor to the MEG signal^37^.

To investigate the effects of cortical excitability on gamma waveform shape we altered the tonic current input to L5 pyramidal and GABAergic interneurons (basket cells). Injection of tonic current may be considered equivalent to changing the excitability of a neuronal population as it raises or lowers the resting potential, bringing it closer or further from firing threshold^34^. We opted to simulate a range of inputs; from -0.5nA to +0.5nA in steps of 0.1nA in the case of pyramidal cells, and from -0.25nA to +0.25nA in steps of 0.1nA in the case of the basket cells. For each condition we simulated a 6-second epoch of gamma, typically resulting in around 300 cycles per condition (total 66 conditions). We performed EMD on the concatenation of all epochs, then performed cycle-by-cycle analysis and waveform shape analysis on simulated data using identical parameters to the MEG data.

Because changing excitability produced large differences in cycle amplitude, statistical comparison on waveform shapes was performed after regressing out overall cycle amplitude as a confound. Average waveform shapes were then calculated for every combination of pyramidal and basket cell excitability and trends calculated using linear regression.

## Results

Participants (N=19) viewed a large, high contrast moving stimulus known to produce strong increases of gamma-band activity and attenuation of alpha/beta-band activity in visual cortex^38–41^ (Fig. 1A), while whole-brain MEG was recorded. Concurrently, short interleaved blocks of anodal, cathodal or sham tDCS were applied to occipital cortex (Oz/Cz montage, Fig. 1B). We chose to interleave short tDCS blocks in order to focus on the effects of tDCS *during* stimulation (‘online’ effects), which are theorised to be driven by changes in resting membrane potential, as distinct from effects *following* stimulation (‘offline’ effects) which may rely on different mechanisms^2,42^. Previous Fourier-based analysis of these data showed no modulation in gamma power, nor of alpha/beta power, by tDCS^23^.

### Intrinsic Mode Functions capture distinct oscillatory responses to moving visual stimulus

After performing Empirical Mode Decomposition to decompose our source reconstructed MEG data into time series of progressively slower neural dynamics, we first checked whether the well-established^38–41^ alpha/beta-band decrease and gamma increase were separated into different Intrinsic Mode Functions (IMFs). Because the grating task has a clear temporal structure, examination of power changes in different epochs (baseline, stimulus on) allowed us to identify expected oscillatory components. Accordingly, we performed conventional time-frequency analysis on each IMF. EMD neatly isolated key oscillatory components into separate modes IMF1-5. IMF1 mainly reflected high-frequency activity that was largely unaffected by task variables (Fig. 3A), IMF2 captured the narrowband gamma increase produced by the onset of the visual stimulus at time 0 (Fig. 3B), IMF3 the beta-band decrease (Fig. 3C), and IMF4 and IMF5 the alpha-band decrease also caused by stimulus onset at time 0 (Fig. 3D,E). Because we were primarily interested in gamma-band activity all subsequent analysis focused on IMF2.

**Figure 4:**
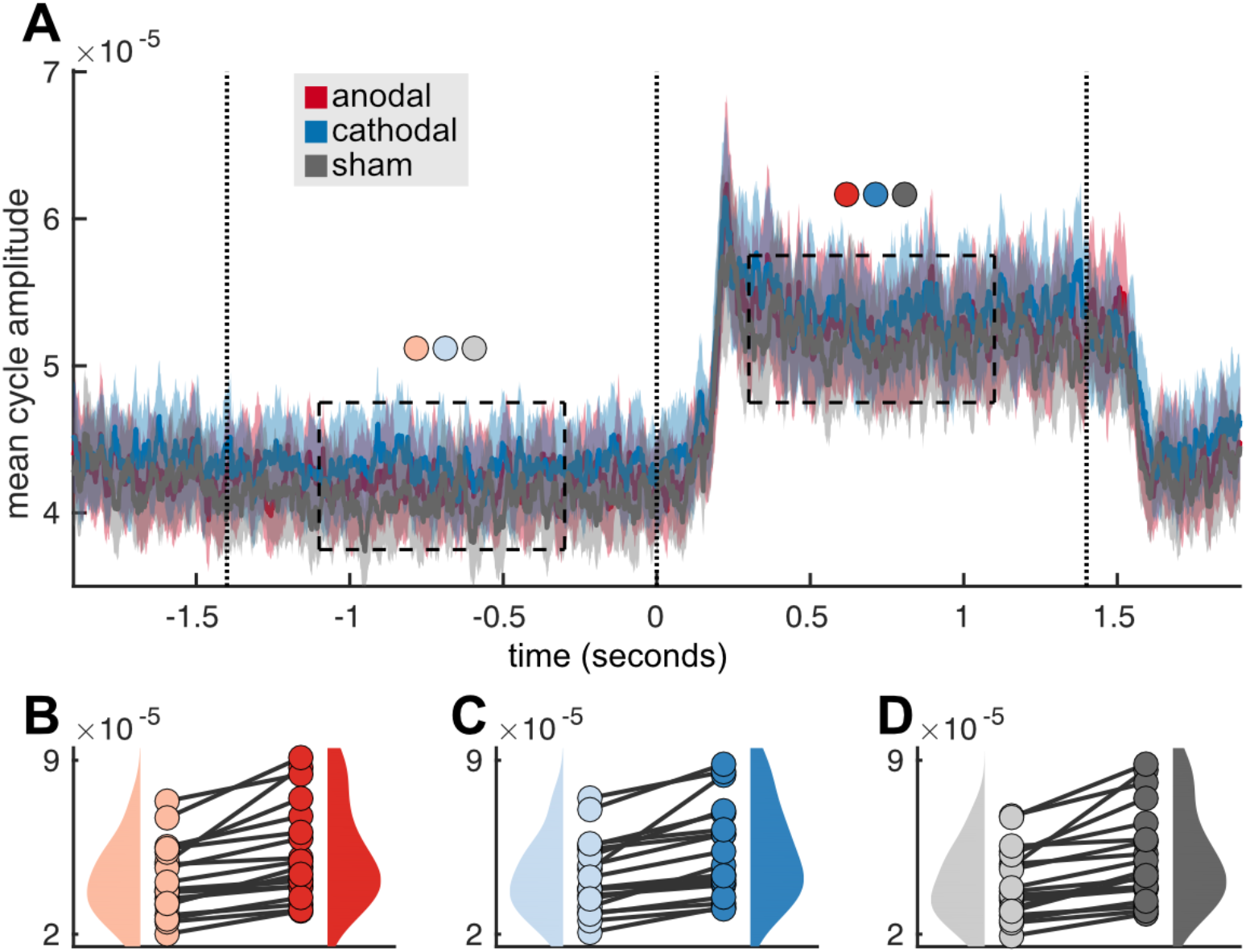
Cycle-wise analysis of gamma mode. A) Mean cycle amplitude across trials for all tDCS conditions (anodal, cathodal, sham). Error bars indicate standard error of the mean. Dashed boxes indicate time windows chosen for statistical analysis. B-D) Individual amplitudes for each participant averaged across the ‘baseline’ and ‘stimulus on’ epochs indicated in A, for Anodal (B), Cathodal (C), and Sham (D) tDCS conditions respectively.

### Cycle-wise metrics confirm tDCS does not alter overall gamma frequency

The EMD toolbox is able to identify individual oscillatory cycles based on analysis of the instantaneous phase of the IMFs^27^. We extracted cycles from IMF2 which contained the stimulus-induced gamma-band response; this resulted in approx 10,000 cycles per participant (mean 9,718, range 5,692-11,821). We then averaged cycle summary information (amplitude, frequency) across trials in each tDCS condition.

In every participant, and every tDCS condition, the onset of the visual grating stimulus caused a strong increase in cycle amplitude (Fig. 3A). Average amplitude increased by 26.1% during anodal tDCS, 25.2% during cathodal tDCS, and 27.4% during sham. Comparing average stimulus-on cycle amplitude with baseline (Fig. 3A, dashed boxes) confirmed strong increases in all tDCS conditions (paired t-tests, Anodal; t(18) = 5.27, p = 5e-5, Cathodal; t(18) = 5.50, p = 3e-5, Sham; t(18) = 5.26, p = 5e-5, Fig. 3B-D).

For further analysis we focused on the ‘stimulus on’ period as this contained the highest-amplitude gamma cycles. We compared mean cycle amplitude and mean cycle frequency between tDCS conditions. No differences were observed in cycle frequency between tDCS conditions (repeated measures ANOVA, F(2,36) = 0.14, p = 0.87). However, a small but significant difference was observed in mean cycle amplitude (repeated measures ANOVA, F(2,36) = 5.98, p = 0.006), driven by a lower amplitude during sham tDCS compared to both anodal tDCS (t(18) = 3.82, p = 0.0013) and cathodal tDCS (t(18) = 3.08, p = 0.0064), which did not differ significantly from each other (t(18) = -1.14, p = 0.27). This is evidence of strong, consistent gamma power increases in the ‘stimulus-on’ time window, which gave us confidence that meaningful comparison of waveform shape in this time window was possible.

### tDCS alters the waveform shape of visual-stimulus-induced gamma oscillations

To determine whether tDCS altered waveform shape of stimulus-induced gamma oscillations we used the EMD toolbox to calculate the instantaneous phase and instantaneous frequency (the temporal derivative of instantaneous phase) of the extracted gamma cycles in the ‘stimulus-on’ time window, then phase-aligned the instantaneous frequencies. Since waveform shape can be re-interpreted as a set of changes in instantaneous frequency across the phase cycle (with a perfect sinusoid having the same instantaneous frequency at all phases and therefore appearing as a horizontal line) transforming cycles into phase-aligned instantaneous frequency allows for meaningful comparison of waveform shapes as a function of task conditions, in our case tDCS (see section, ‘The logic of cycle phase alignment’, Fig. S1 for a conceptual sketch, and for full implementational details see^26^).

Focusing on the gamma-band IMF, and the visual stimulation epoch where strong gamma-band activity was observed, we found that waveform shapes significantly differed between anodal and cathodal tDCS conditions (Fig. 5A, cluster-based permutation test, p = 0.0138). Visual inspection revealed maximal differences in the second half of the phase cycle (p < 0.05 uncorrected at every phase point from 53%-99% of the cycle). This means that tDCS mainly altered the shape of *one half-wave* of the cycle, while not altering the other half-wave. During anodal tDCS, cycles had slower instantaneous frequency in the descending half-wave relative to cathodal stimulation. In other words, after the down-going zero crossing, gamma cycles under anodal tDCS took relatively longer, and under cathodal tDCS relatively shorter, to return to their initial state.

**Figure 5:**
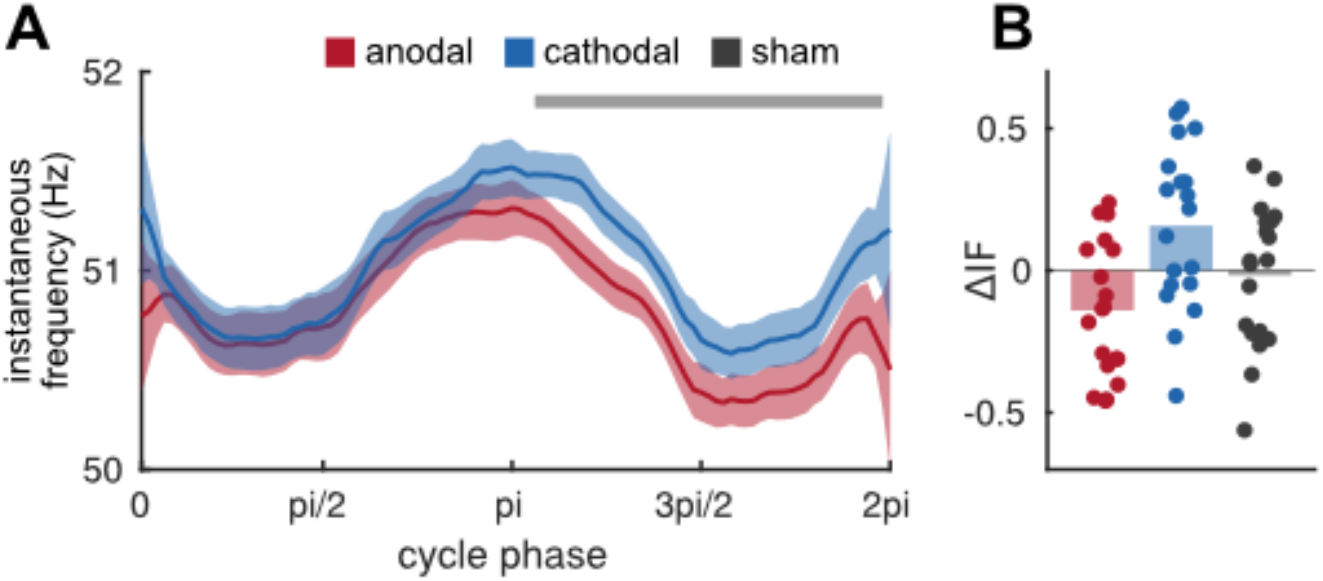
tDCS affects gamma waveform shape. A) Phase-aligned instantaneous frequencies of gamma IMF during ‘visual stimulus on’ period, for anodal (red) and cathodal (blue) stimulation. Shaded areas indicate standard error of the difference. Grey horizontal line indicates time points of significant difference at p < 0.05 uncorrected (overall significant difference determined by cluster-based permutation test). 0, pi and 2pi represent zero crossings, pi/2 and 3pi/2 represent peaks/troughs (note that ‘peak’ and ‘trough’ are inherently ambiguous, see section ‘Resolving Dipole Sign Ambiguity’). B) Mean change in instantaneous frequency over window of significant anodal-cathodal difference shown in A, for each tDCS condition, relative to the mean of all three conditions.

We next determined whether gamma waveform shape in either tDCS polarity differed from sham, by averaging instantaneous frequency across the timepoints from the cluster test and comparing all three conditions (Fig. 5B). Qualitatively, mean gamma instantaneous frequency appeared to be intermediate between anodal and cathodal. However, statistical testing revealed that cathodal instantaneous frequency was not significantly greater than sham (t(18) = 1.59, p = 0.13) and anodal not significantly smaller than sham (t(18) = 1.35, p = 0.19). Note that anodal and cathodal did differ significantly (t(18) = 2.80, p = 0.0117); however, this difference is trivial as the window selected for testing was based on a significant anodal-cathodal difference identified in the cluster permutation test (‘double-dipping’).

Because instantaneous phase cannot be accurately estimated when amplitude is low, we excluded the 30% of cycles with lowest amplitude for each participant. Control analyses using a range of amplitude cut-offs revealed that the specific cut-off choice did not impact the results (Supplementary Materials, Fig. S2).

The above is strong evidence that online tDCS to occipital cortex did impact visual stimulus-induced gamma activity, that – rather than altering gamma amplitude and frequency as initially hypothesized^23^, tDCS altered gamma *waveform shape*, and that the shape alterations were confined to one half of the oscillatory cycle.

### Spiking network model predicts halfwave-specific shape alterations

tDCS, particularly applied *online* as in the present study, is thought to act on resting membrane potential^8^. However, the relationship between resting potential and population-level phenomena such as gamma oscillations is complex. One possibility to elucidate this relationship is to derive predictions from simulation. To determine whether the half-wave-specific effects we observed in our MEG data were biophysically plausible, and to indicate possible mechanisms, we used the Human Neurocortical Neurosolver^34^ (HNN) – a computational neural model that simulates the electrical activity of neocortical circuits – to simulate the effects of altering resting membrane potential on a gamma-generating neuronal population.

HNN can implement a PING-type model of gamma oscillations whereby a Poisson input to excitatory cells causes excitatory spiking that activates inhibitory cells, producing a rhythm proportional to the excitation strength and inhibitory time constant^16^. The HNN documentation refers to this model as ‘weak ping’^34^. We adapted this model by adding a range of weak positive or negative tonic inputs to the somata of layer 5 pyramidal and basket cells. For full details see ‘Methods’.

The simulated HNN network generated robust gamma oscillations in all conditions (Fig. 6A). We then extracted cycle information and applied waveform shape analysis to the synthetic gamma cycles exactly as previously applied to the MEG data.

**Figure 6:**
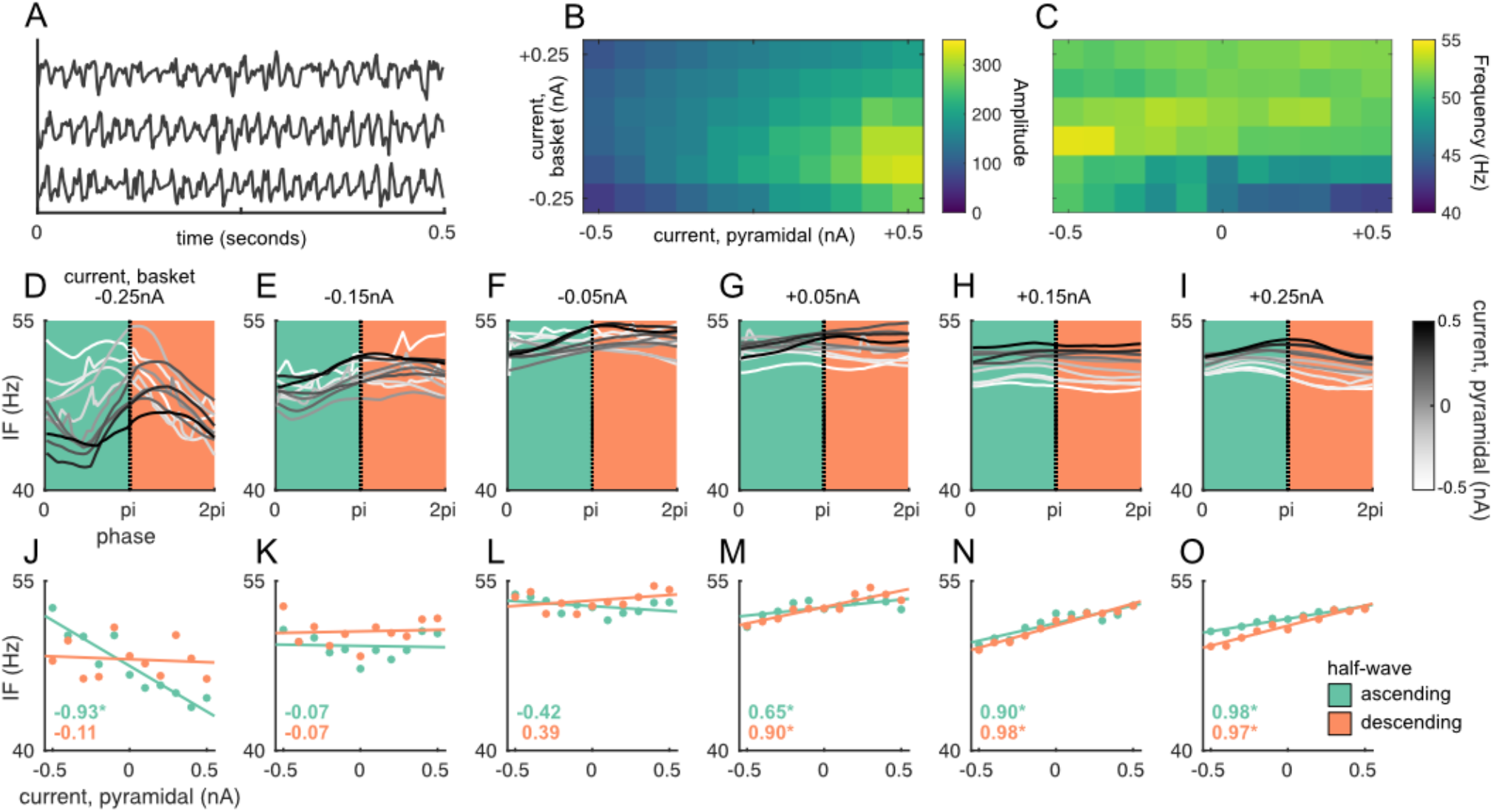
Waveform shape analysis of simulated gamma oscillations. A) Representative epochs of simulated gamma oscillations under different excitability changes to pyramidal and basket cells. B) Average cycle amplitude as a function of different current levels to pyramidal (rows) and basket cells (columns). C) as B, but for cycle frequency. D-I) Average gamma waveform shapes split up by excitability level of basket cells (separate plots). Line colour (white-black) indicates pyramidal excitability. Background colour (green/orange) indicates half-wave; green-ascending, orange = descending. J-O) Instantaneous frequency averaged over ascending (green dots) and descending (orange dots) halfwaves. Lines indicate linear fits. Inset numbers indicate r-values associated with linear fits

When examining overall cycle characteristics (amplitude and frequency), altering tonic inputs to pyramidal cells in the simulated network primarily affected cycle amplitude (Fig. 6B). At all levels of basket cell excitability, more current to pyramidal cells increased cycle amplitude. Effects on cycle frequency were relatively small (Fig. 6C); primarily, when negative currents were applied to basket cells there was a decrease in cycle frequency with more current to pyramidal cells, but this effect was not observed at higher amplitudes.

We then performed waveform shape analysis for all cycles by redescribing the cycle as phase-aligned instantaneous frequency, identically to the MEG data. Here a highly interesting pattern of results was observed: When basket cell excitability was lowest, altering pyramidal cortex excitability appeared to produce changes in mean waveform shape that were confined to one half-wave (Fig. 5D, green section), and absent in the other half-wave (orange section). Correlation analysis confirmed the presence of a significant linear trend (r = -0.93, p = 0.00003) in the ascending half-wave that was absent in the descending half-wave (r = -0.11, p = 0.75). At higher basket cell excitability this effect was absent: Weakly negative current to basket cells led to the absence of any effect of pyramidal current on waveform shape in either halfwave (Fig. 5E,F,K,L). Positive current to basket cells led to effects of pyramidal excitability on waveform shape; more current to pyramidal cells increased instantaneous frequency, but in both half-waves in the same direction (Fig. 5G-I,M-O). In other words, here pyramidal current had a positive effect on *overall cycle frequency*, rather than changing waveform shape.

The above results indicate that the relationship between cell excitability and waveform shape is complex and nonlinear. However, they support the notion that halfwave-specific alterations to gamma waveform shape by cortical excitability, as we observed in our MEG data, are biophysically plausible, that pyramidal cells are the likely locus of those effects, and that this pattern of effects is more likely to arise when basket cells are in a state of relatively low excitability.

## Discussion

Via a combination of theoretical and experimental approaches, we investigated the effects of altering resting membrane potential on gamma oscillations. By reanalysing a dataset where tDCS was applied to the occipital cortex while participants observed visual stimuli known to increase gamma activity^38–41^, we showed that – even though previous analysis^23^ indicated that tDCS did not alter gamma amplitude and frequency, as measured using conventional time-frequency analysis^23^ – here we show that it did produce robust, specific changes in the *waveform shape* of gamma activity. Specifically, gamma waveform shape was relatively compressed under cathodal tDCS and relatively expanded under anodal tDCS, with these changes confined to one half-wave of the gamma cycles, and absent in the other half-wave. This finding was robust to control analyses using different amplitude cut-offs.

Furthermore, we confirmed the biophysical plausibility of halfwave-specific alterations of waveform shape, by simulating gamma-generating neuronal networks with differing degrees of cortical excitability. Specifically, when inhibitory basket cells were in a state of relatively low excitability, we observed that pyramidal cell excitability robustly and parametrically modulated waveform shape in one halfwave only. This indicates that the observed MEG effects likely resulted from alteration of the membrane potential of pyramidal cells. This is consistent with the finding that – due to their morphology – layer 5 pyramidal neurons are most likely to be sensitive to polarisation by sub-threshold electric fields^28^. The convergence of theoretical and experimental results increases confidence that the findings arise from a true physiological effect and provides new insights on the effects of tDCS.

It is noteworthy that, while both experimental and simulation findings showed halfwave-specific alterations of gamma waveform shape – the differences in the MEG data were observed in the *descending* halfwave (from pi to 2pi) and the differences in the simulation were observed in the *ascending* halfwave (from 0 to pi). Importantly, and as discussed, the sign of the source-reconstructed MEG data is arbitrary, so strong conclusions cannot be drawn about whether a given halfwave corresponds to the peak or trough of an oscillation. We used a group-PCA approach (section ‘Resolving dipole sign ambiguity’) to ensure sign consistency *across participants*. While this procedure allows meaningful group statistics and comparisons, it cannot resolve the ‘correct’ sign with respect to any ground truth. Nevertheless, in both data and model the observed pattern was similar; altering cortical excitability altered waveform shape in one half-wave, while preserving it in the other half-wave.

This study highlights the importance of waveform shape as an additional feature of neuronal oscillations^43^. Much attention has been focused on transcranially altering oscillatory activity; many studies seek to ‘entrain’ neuronal oscillations by applying rhythmic transcranial stimulation, typically tACS^44^ or rTMS^45^. The intention here is typically to change oscillatory *amplitude*. Our approach can be considered complementary: Here we show that the application of DC fields can also alter oscillatory activity along another dimension – the shape of the oscillation – by altering biophysical properties of the neurons that interact to produce an oscillation in the population-level signal.

Our findings are interesting in the context of the ‘communication through coherence’ hypothesis, that posits gamma synchronisation as a fundamental mechanism of inter-areal neuronal communication^11^. Communication through coherence relies on the alignment of excitatory windows, ensuring a spike arriving from an upstream region arrives at a downstream neuron in a period where that region is maximally excitable, as opposed to a phase of relatively low excitability. Gamma-band synchronisation can be understood as a mechanism to maximise the efficacy of upstream output on a downstream region. In this context, alterations of gamma waveform shape are particularly interesting; they may indicate a relative lengthening or shortening of the population ‘excitatory window’. This, in turn, may have implications for improving or disrupting inter-areal communication.

A limitation of the current study was that our simulations focused solely on cortical layer 5. *In Vitro* experimental work suggests that layer 5 pyramidal cells are most strongly affected by subthreshold electrical stimulation^28^, possibly due to their morphology. Furthermore, theoretical modelling work suggests that – due to their longer apical dendrites – gamma rhythms produced in layer 5 are likely to dominate M/EEG signals measured outside the head^37^. However laminar recordings from monkey V1 suggest that visual gamma is likely generated in superficial and granular layers, with relatively less contribution from layer 5^36,46^, gamma spike-field coherence is also stronger in superficial than deep layers^35^, and simultaneous EEG-fMRI suggests variability in gamma in the EEG signal correlates most strongly with BOLD in superficial layers^47^. It might even be that multiple gamma rhythms with different laminar origins contribute independently to the MEG signal^37^. Future modelling work should therefore consider both contributions from other layers as well as possible inter-laminar interactions.

Finally, we believe the combination of computational simulations and neuroimaging experiments could in general be highly illuminating to further investigate the mechanisms of tDCS. A great deal of interesting work has been done showing that stimulating a particular brain region changes some aspect of cognition and behaviour. However, it should be noted that the effects of tDCS on behaviour and ‘higher-order’ cognition have been characterised as somewhat weak and inconsistent^6,7^. Since the mechanism of action of tDCS – changing cortical excitability – is best understood at the neuronal level, a useful approach to reduce this inconsistency would be to ‘bridge the gap’ between neurons and behaviour by combining brain stimulation, neuroimaging and computational modelling. Neural population models like HNN enable quantitative predictions about how altering cortical excitability will change activity at the neuronal population level. This generates hypotheses about population activity that can be tested with neuro-imaging methods such as MEG, that are sensitive to population signals. Such a three-pronged ‘computational neurostimulation’^48,49^ approach could revolutionise brain stimulation and bring us closer to applications for cognitive and clinical benefit.

## Acknowledgements

The authors thank Sophie Esterer and Jim Herring for help with experimental design and MEG data collection.

## Author Contributions

TRM, OJ and TOB planned the study. TRM acquired the data. TRM analysed the data. AJQ contributed analysis tools and code. TRM wrote the manuscript. All authors revised and edited the manuscript.

## Supplementary Materials

### The logic of cycle phase alignment

NB: This conceptual summary glosses over important implementational detail. A full explanation can be found in^26^.

**Figure S1:**
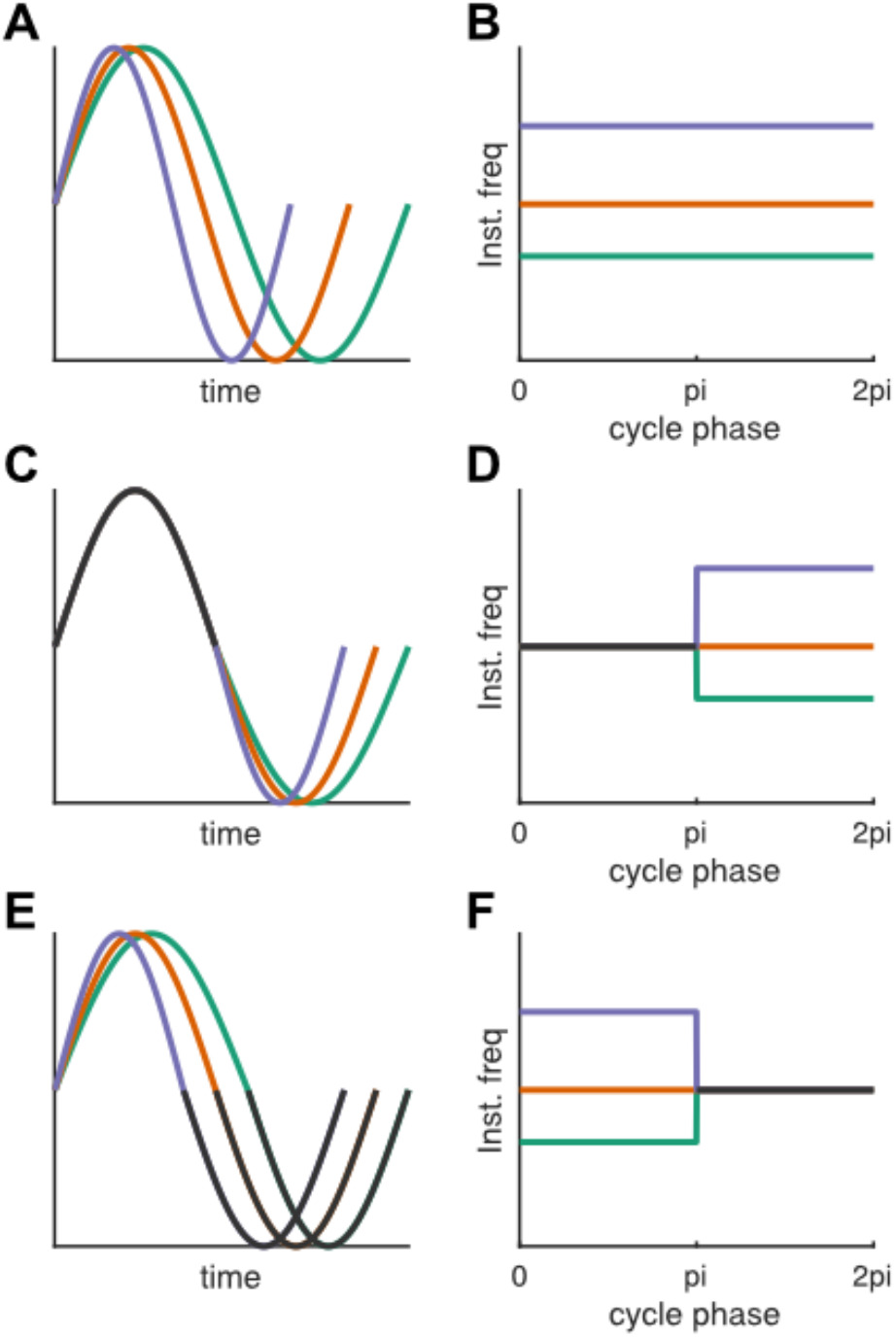
Explanation of phase-alignment logic. A) Perfect sinusoids with different frequencies. B) Phase-aligned instantaneous frequency of the three waveforms in A. C,E) Complex waveforms which differ in part of the phase cycle. D,F) Phase-aligned instantaneous frequency of the three waveforms in C,E.

All cyclical phenomena can be re-conceived in terms of their instantaneous frequency at different points in their phase cycle. Fig. S1A shows three simple sinusoidal waveforms with different frequencies. Comparing the waveform shapes in the time domain is difficult because the cycles do not have the same duration; the fast, purple cycle has finished before the slow, green cycle reaches its trough. However, when re-describing them as phase-aligned instantaneous frequency (Fig. S1B) their similarity becomes obvious. Because sinusoids have unchanging instantaneous frequency across the phase cycle each appears as a horizonal line at uniform instantaneous frequency, with the fast purple cycle having the highest frequency. Using this phase-alignment approach, different portions of the cycle can be reasonably compared despite unequal durations. Fig S1C shows three complex waveforms. All waveforms have the same ascending halfwave (black trace), but differ in the speed of the descending halfwave (purple = fast, orange = intermediate, green = slow). Again, re-expression in terms of phase-aligned instantaneous frequency (Fig. S1D) simplifies the process of identifying exactly where and how in the cycle the waveforms differ. Fig. S1E,F reverses the situation in Fig. S1C,D; the three cycles have different ascending half-waves but identical descending half-waves. Here the misalignment of the descending half-waves (Fig. S1E) makes comparison in the time-domain complicated. However, aligning the cycles in the phase domain (Fig. S1F) allows simple comparisons of where they are similar and where they are different.

### Waveform effects do not depend on choice of amplitude cutoff

Determining waveform shapes relies on estimating instantaneous phase, which cannot be estimated accurately if the cycle has a low amplitude. Therefore, in our analysis of gamma waveform shape (main Fig. 5) we excluded from analysis all cycles with a mean instantaneous amplitude below 30% of the mean for that IMF and participant. We wanted to make sure that the pattern of observed results did not depend on this arbitrary choice of threshold. Therefore, for illustration, we repeated the analysis at a range of cutoffs from 0% (all cycles included) to 95% (only highest-amplitude cycles included). Qualitatively, the observed pattern – a difference between gamma waveform shape between anodal and cathodal stimulation, in the descending half-wave (from pi to 2pi) – was observable for a wide range of cutoffs from 20-60%. We suspect that below a 20% cutoff low-amplitude cycles distort the mean waveform shapes, and above 60% we lack sufficient SNR to estimate the shapes due to insufficient numbers of cycles.

**Figure S2:**
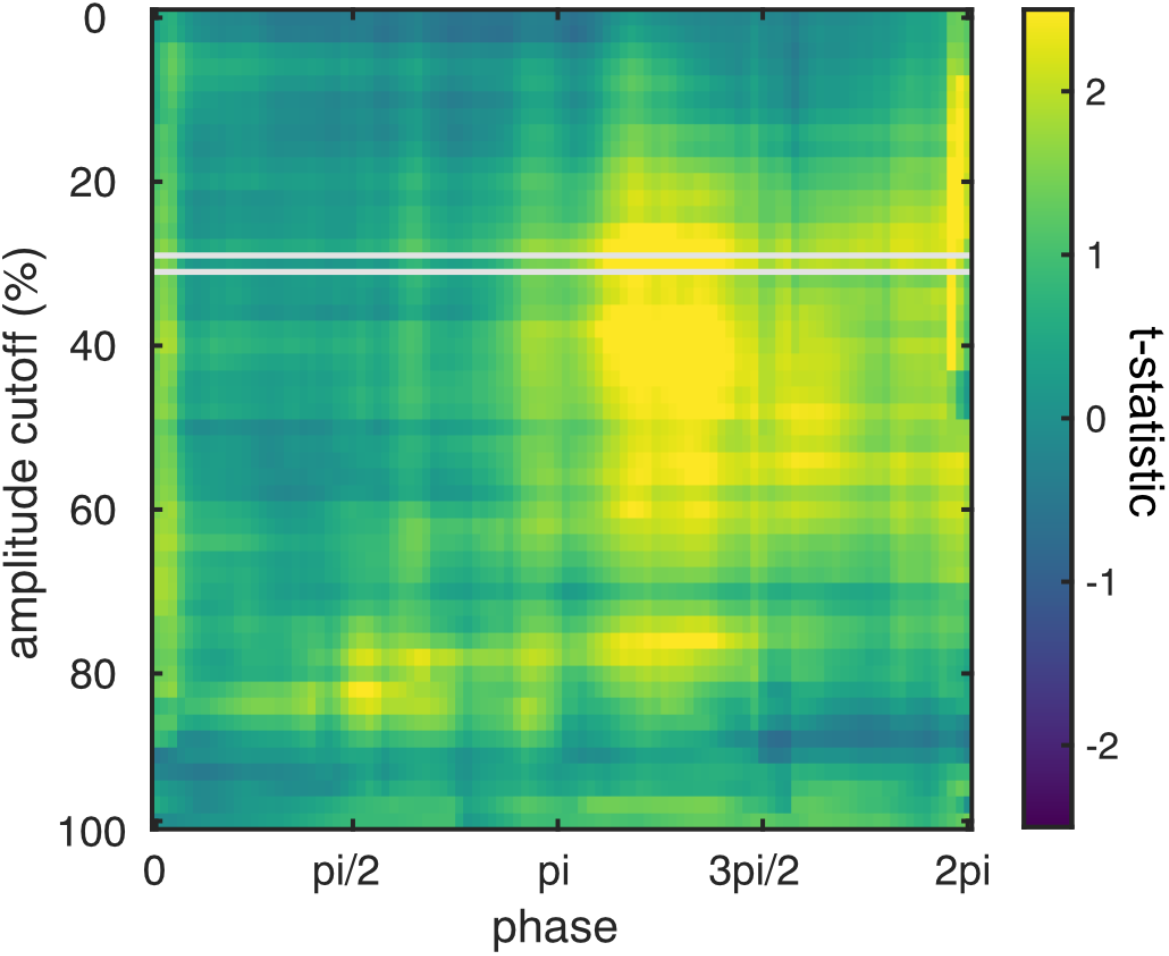
Statistical comparison of gamma waveform shape between anodal and cathodal tDCS, for a range of amplitude cutoffs. Grey box indicates the amplitude cutoff chosen for the main analysis (fig5).

